# Widespread coexistence of self-compatible and self-incompatible phenotypes in a diallelic self-incompatibility system in *Ligustrum vulgare* (Oleaceae)

**DOI:** 10.1101/2020.03.26.009399

**Authors:** Isabelle De Cauwer, Philippe Vernet, Sylvain Billiard, Cécile Godé, Angélique Bourceaux, Chloé Ponitzki, Pierre Saumitou-Laprade

## Abstract

The breakdown of self-incompatibility (SI) in angiosperms is one of the most commonly observed evolutionary transitions. While multiple examples of SI breakdown have been documented in natural populations, there is strikingly little evidence of stable within-population polymorphism with both inbreeding (self-compatible) and outcrossing (self-incompatible) individuals. This absence of mating system polymorphism corroborates theoretical expectations that predict that in/outbreeding polymorphism is possible only under very restricted conditions. However, theory also predicts that a diallelic sporophytic SI system facilitates the maintenance of such polymorphism. We tested this prediction by studying the mating system of *Ligustrum vulgare* L., an entomophilous hermaphroditic species of the Oleaceae family. Using stigma tests with controlled pollination and paternity assignment of open-pollinated progenies, we confirmed the existence of two self-incompatibility groups in this species. We also demonstrated the existence of self-compatible individuals in different populations of Western Europe arising from a mutation affecting the expression of the pollen component of SI. We then estimated the selfing rate in a garden experiment. Our results finally show that the observed low frequency of self-compatible individuals in natural populations is compatible with theoretical predictions only if inbreeding depression is very high.

## Introduction

Mating system transitions are among the most frequent evolutionary transitions in a wide range of taxa including angiosperms, fungi, algae, and bryophytes (Stebbins 1957; Billiard et al. 2011). In particular, the loss of self-incompatibility (SI) in flowering plants has occurred many times independently, even within families (Goodwillie 1999; Igic et al. 2008; Goldberg et al. 2010). Theoretical work has shown that SI can be lost or maintained depending on the balance between the opposing effects of a self-compatible (SC) mutation on female fitness (inbreeding depression vs. pollen limitation), male fitness (siring success associated with mate availability vs. pollen discounting), and its own direct fitness advantage (Fisher 1941; Charlesworth and Charlesworth 1979; Porcher and Lande 2005a; Gervais et al. 2014; Van de Paer et al. 2015).

In addition to predicting the conditions under which SI can be maintained or lost, models also predict the stable coexistence of SC mutants with a set of functional SI alleles, but only under a restricted set of conditions (Charlesworth and Charlesworth 1979; Porcher and Lande 2005b; Van de Paer et al. 2015). First, the strength of inbreeding depression should be intermediate, because high values would result in an exclusion of SC mutants from the population, while low values would lead to a rapid fixation of a SC phenotype. Second, the number of functional self-incompatibility alleles (S-alleles) should be limited otherwise the reproductive advantage of SC genotypes relative to SI ones would be offset, resulting in the exclusion of SC mutants. When both conditions are fulfilled, the stable maintenance of the SI/SC polymorphism within a population is possible without being affected by the genetic architecture underlying the SC phenotype (*i*.*e*. the causal mutation can either affect the self-incompatibility locus itself or an unlinked locus that interacts epistatically with the self-incompatibility locus, Charlesworth and Charlesworth 1979; Porcher and Lande 2005b). Likewise, the SI/SC polymorphism can be maintained whether the mutation affects the male or female function only (Uyenoyama et al. 2001; Gervais et al. 2011; Van de Paer et al. 2015). Finally, the stability of the SI/SC polymorphism is not affected by the characteristics of the inbreeding depression, which can be fixed or variable, and due to lethal recessive or mildly deleterious mutations (Gould and Vrba 1982; Porcher and Lande 2005b). Overall, the crucial parameters for a stable maintenance of the SI/SC polymorphism are thus the strength of inbreeding depression and the number of functional S-alleles. It should be noted that this second condition should rarely be met in natural population, where the number of functional S-alleles is typically very large (Castric and Vekemans 2004) and SI/SC polymorphisms are thus predicted to be rare or transient, which is corroborated by observations.

To the best of our knowledge, the empirical evidence of stable SI/SC polymorphism is indeed scarce and ambiguous in natural populations. For instance, natural populations of *Laevenworthia alabamica* have been shown to be either SI or SC, with no evidence of mixed populations (Lloyd 1965; Busch and Schoen 2008). Similarly, a set of wild populations of *Arabidopsis lyrata* sampled in North America proved to be either predominantly SI or SC, except one population where SI and SC individuals were equifrequent (Mable et al. 2017). Hence, even though *L. alabamica* and *A. lyrata* show SC/SI polymorphism, [SC] and [SI] phenotypes are generally spatially segregated. This segregation suggests either that the SC/SI polymorphism is only transient with SI ultimately being lost as suggested in North American *A. lyrata* populations (Mable et al. 2017), or, if stable, the observed polymorphism is a direct consequence of metapopulation dynamics with *L. alabamica* SC populations being found at the species’ range margin (Lloyd 1965).

Overall, theoretical expectations and empirical observations in multi-allelic SI species seem to coincide. If so, then theoretical predictions should also be correct regarding the existence of a SC/SI polymorphism in the particular case of homomorphic diallelic self-incompatibility systems (DSI). For example, Van de Paer et al. (2015) showed that, in the absence of unisexual individuals, [SC] phenotypes are expected to invade a SI population with only two S-alleles and three stable situations are possible (Van de Paer et al. 2015): either SI is lost as the [SC] phenotype invades and goes to fixation, or the [SC] phenotype stably coexists with one or two [SI] phenotypes. The common privet (*Ligustrum vulgare*), an Oleaceae species, is an ideal species to test these theoretical predictions, which is the main goal of this paper.

The common privet is described as a purely hermaphroditic species (*i*.*e*. it does not show unisexual individuals that would preclude the invasion of mutant [SC] phenotypes, Van de Paer et al. 2015) and, even though nothing is known about its mating system, it is strongly suspected to have a DSI like all other Oleaceae species that have been studied so far. To date, the occurrence of DSI has been confirmed in *Fraxinus ornus* (Vernet et al. 2016), *F. excelsior* (Saumitou-Laprade et al. 2018), *Phillyrea angustifolia* (Saumitou-Laprade et al. 2010), *Olea europaea* subsp. *europaea* (Saumitou-Laprade et al. 2017a) and *O. europaea* subsp. *laperrinei* (Besnard et al. 2020). The first goal of this work was thus to test whether the common privet expresses DSI. If the number of S-alleles is indeed limited to two in wild *L. vulgare* populations, either SI is expected to have been lost because the [SC] phenotype has invaded and gone to fixation, or the [SC] phenotype stably coexists with one or two [SI] phenotypes as theoretically expected (Van de Paer et al. 2015). Thus, the second goal of this work was to determine whether [SC] phenotypes can indeed be found in the study species *in natura*. To do so, we screened a set of wild populations for SI/SC behavior using stigma tests. We also transferred a set of these individuals to an experimental garden and performed controlled stigma tests in a diallelic crossing design in order to assess compatibility relationships among [SI] and [SC] individuals used as pollen recipients and pollen donors. Third, because Van de Paer al. (2015) predict a stable SC/SI polymorphism in species with DSI only if selfing rates are high, we conducted a paternity analysis in the experimental population to estimate the rate of self-fertilization in open-pollinated maternal progenies. Finally, we applied the model developed by Van de Paer et al. (2015) to our experimental results to formulate quantitative predictions on the level of inbreeding depression required to maintain the mating system observed in *L. vulgare* in natural populations.

## Materials and methods

### Plant material and study populations

#### Ligustrum vulgare *in the Oleaceae family*

*L. vulgare* (common privet or European privet) is hermaphroditic. It is a deciduous or semi-evergreen shrub, native to Europe, North Africa, and western Asia (Fig. 1). It has been introduced from Europe to North America where it is considered invasive. One individual plant can produce more than 10 000 fruits (Obeso and Grubb 1993) and seed dispersal can be facilitated by birds or animals during winter. Individuals can also reproduce vegetatively. Like the majority of species in the Oleaceae family, *L. vulgare* is an insect-pollinated species with petaliferous, fragrant and nectariferous flowers.

**Figure 1:**
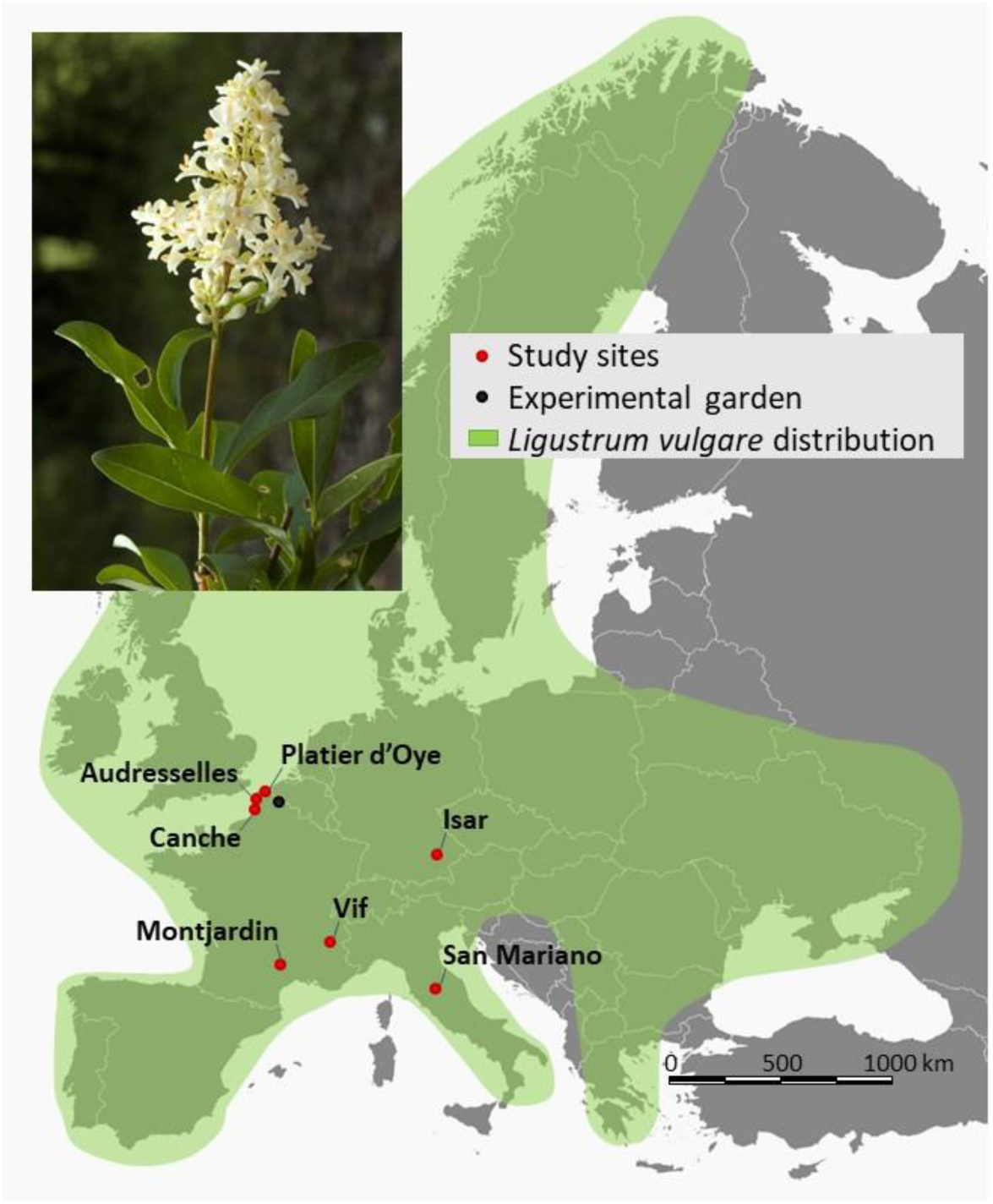
*Ligustrum vulgare* geographic species range across Europe and position of the seven studied natural populations. Photo credit: Marijke Verhagen, Saxifraga Foundation.

#### Identification of tester genotypes

To verify the occurrence of DSI in *L. vulgare*, the first step was to identify two cross-compatible genotypes whose pollen would then be used as tester pollen across a set of wild populations. If the study species indeed possesses only two SI groups, then any individual used as a pollen recipient would be compatible with only one of these tester genotypes. Compatibility was assessed using stigma tests, following Saumitou-Laprade et al. (2017a), on six individuals in a study site close to the laboratory facilities (Marais de Péronne, N 50°33’30.298”, E 3°9’59.778”). When the pollen recipient and the pollen donor are compatible, several pollen tubes converge and grow through the stigmatic tissue towards the style and then through the style towards the ovules (*e*.*g*. Fig. 2 panels A4, A5, B4, and B5). The absence of pollen tubes (or only short pollen tubes that do not reach the style) was used as the criterion to score incompatibility (*e*.*g*. Fig. 2 panels A1, A2, B1, and B2).

**Figure 2:**
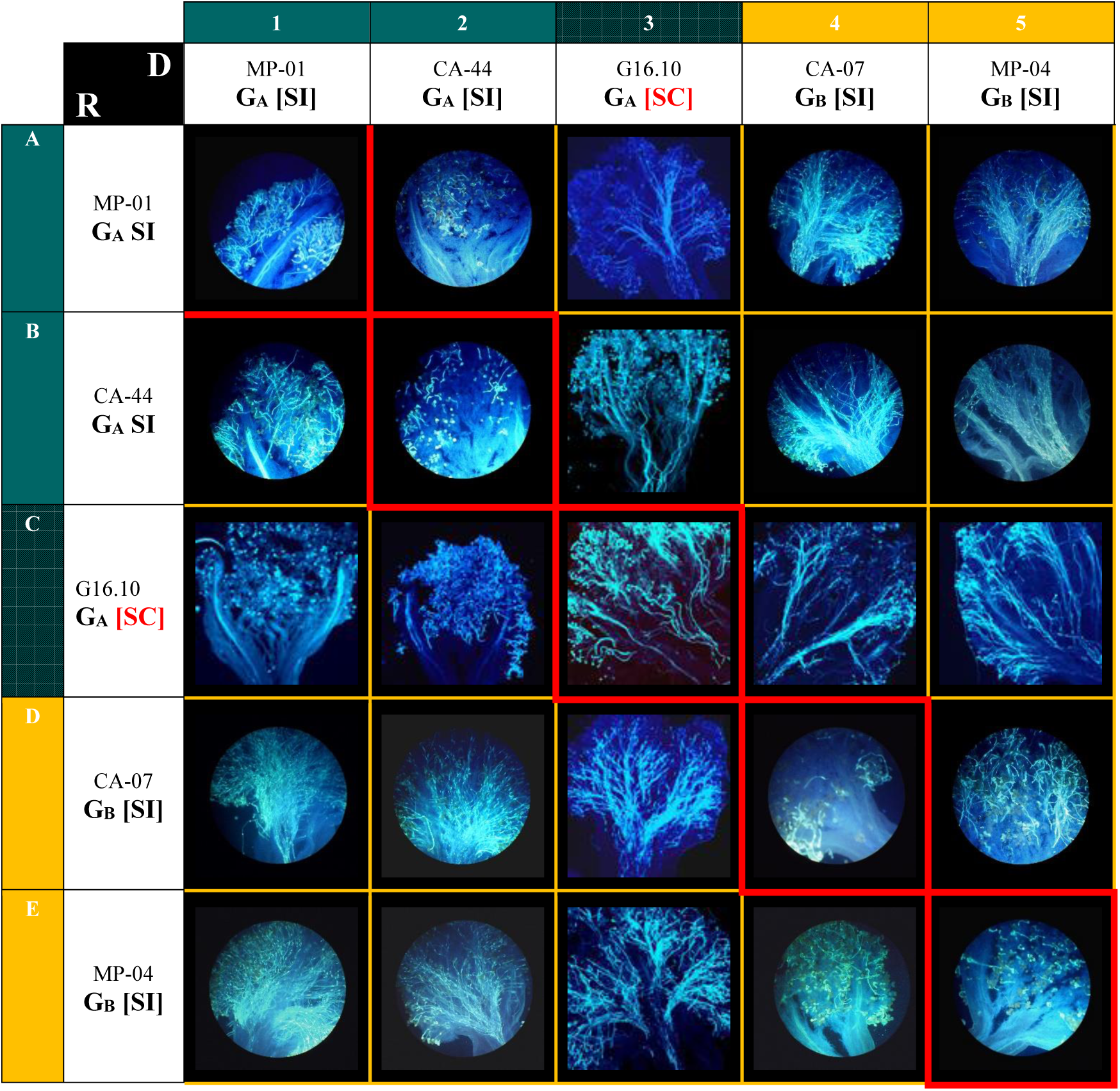
Illustration of the different compatibility relationships observed using stigma tests in *Ligustrum vulgare*. In the diallelic-design cross, plants where used as pollen donors (D) and as pollen recipients (R). Each column (1 to 5) corresponds to a genotype used as the pollen donor on five recipient genotypes (rows A to E). Panels outlined in red correspond to self-pollinations. G_A_[SI] and G_B_[SI]: self-incompatible genotype belonging to the self-incompatibility groups A and B respectively; G_A_[SC] self-compatible genotype which stigmas are incompatible with pollen from G_A_ genotypes.

Once a cross-compatible tester pair (MP-01 and MP-04) was identified, freshly collected inflorescences containing a few open flowers from these two genotypes were transferred to the laboratory. Open flowers were eliminated and inflorescences were then placed in controlled conditions until flower buds opened. The four-lobed corolla with attached anthers were collected with forceps and stored in a Petri dish. Dehiscence was then induced by placing corollas with anthers in a dry atmosphere for 4 h. Corolla with dehisced anthers were immediately stored at -80°C.

#### Screening wild populations for DSI and [SI]/[SC] phenotypes

Pollen from these two cross-compatible tester genotypes was used to assess compatibility on 184 individuals sampled in seven European natural populations located in five geographically distinct regions in France (North Sea coast, Cevennes, Alps), Germany (Bavaria) and Italy (Umbria) (see Fig. 1 and Table 1). Any individual incompatible with MP-01 and compatible with MP-04 was classified as belonging to an incompatibility group called “G_A_”. Conversely, individuals compatible with MP-01 and incompatible with MP-04 were classified as “G_B_”. If more than two compatibility groups occur in this species, some individuals would be compatible with both testers, forming a potential “G_N_” group. In addition to these DSI tests, a subsample of trees (142 individuals) was also tested for self-compatibility to identify potential [SC] phenotypes (Table 1). Once individuals showing a [SC] phenotype were identified, we estimated the [SC] phenotype frequencies in each population by applying simultaneous confidence intervals for multinomial proportions according to the method described in Sison and Glaz (1995), using the MultinomCI function in the DescTools package in R (Signorell 2019). For further tests, 236 individuals (plant cuttings or seeds) from six of these wild populations were transferred to the experimental garden at the University of Lille (Fig. S1). Most of them (88%) were also assessed for SI and assigned to a SI group, allowing us to identify two [SC] individuals.

**Table 1:**
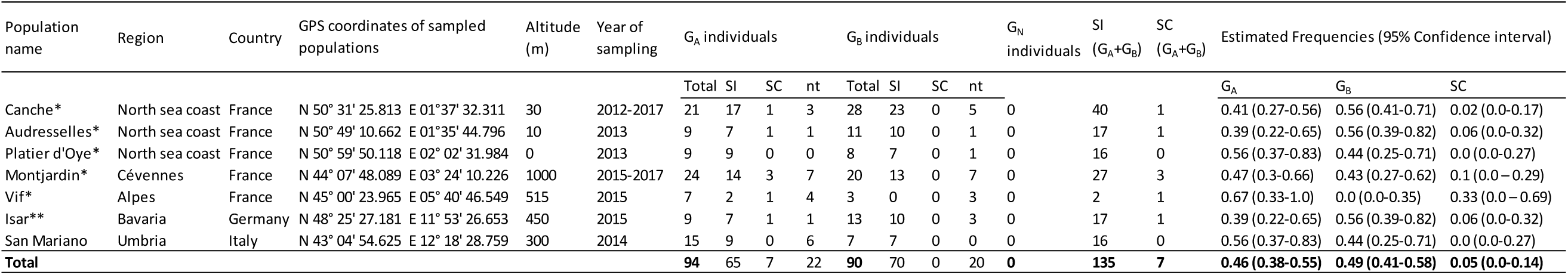
Screening of natural population of *Ligustrum vulgare* for incompatibility groups using stigma tests. Sampled individuals were tested as pollen recipients using the pollen from two inter-compatible tester plants (MP-01 and MP-04). G_A_: individuals incompatible with MP-01 and compatible with MP-04; G_B_: individuals compatible with MP-01 and incompatible with MP-04; G_N_: individuals compatible with both MP-01 and MP-04. SI: individuals for which self-pollen did not germinate on their own stigmas; SC: individuals for which self-pollen germinated on their own stigmas. *Populations from which cuttings of individual genotypes were transferred to the experimental garden; **Population from which collected seeds were sown and grown in the experimental garden.

#### Pre- and postzygotic analysis of cross-compatibility behavior among [SI] and [SC] *L. vulgare*

- **Prezygotic analysis:** Stigma tests performed in a diallelic design To precisely describe the cross-compatibility behavior of [SC] individuals detected in the population screening, we performed controlled pollinations in a full diallelic crossing scheme with four G_A_ (CA-44, MP-01, VIF-09, and VIF-16), four G_B_ (CA-07, CA-31, G13-08, and MP-04), and two SC genotypes (G16-10 and MJ-05). In this reciprocal diallele design, each genotype was tested as a pollen donor and pollen recipient for cross-compatibility behavior with all other genotypes using the stigma test (as performed in *Olea europaea* subsp. *europaea*: (Saumitou-Laprade et al. 2017a; Saumitou-Laprade et al. 2017b). Each of these 100 crosses was replicated on three flowers. We then scored the germination (or the absence of germination) of pollen tubes in stigmas for the 3×100 crosses.
- **Postzygotic analysis:** paternity success and selfing assessed in a genetically isolated experimental population

Paternity analyses were then performed to confirm the results of the stigma tests at the postzygotic stage. In 2017, 10 individuals, including 9 of the genotypes used for the prezygotic diallelic stigma test, produced substantial amounts of seeds in open pollination. These plants included three [G_A_] (CA-44, MP-01 and VIF-09), five [G_B_], (CA-07, CA-31, G13-08, MP-04 and MJ-06-1), and two [SC] genotypes (G16-10 and MJ-05). Progenies were collected in autumn and divided into two groups. In the first group, offspring were genotyped at the embryo stage (498 offspring, with 50 embryos per mother plant, except for MP-04 for which only 48 seeds were available). In the second group, seeds from six progenies (two [G_A_]: CA-44 and VIF-09; two [G_B_]: CA-31 and G13-08; and the two [SC]: G16-10 and MJ-05) were germinated in the greenhouse and grown until each seedling had several leaves before genotyping (711 offspring collected on six mother plants, with an average of 118 seeds per mother plant).

##### DNA extraction and genotyping

Total genomic DNA from the 236 adults in the experimental plot and the 1209 offspring (498 embryos and 711 seedlings) was extracted and purified using the NucloSpin 96 Plant II Kit (Macherey Nagel, Duren, Germany) following the manufacturer’s protocol. For adults and seedlings, DNA was extracted from 10 to 20 mg of lyophilized leaves. For embryos, the full embryo was separated from the endosperm before lyophilization and DNA extraction. Five polymorphic microsatellite loci were amplified in a single multiplex reaction (Supplementary Table 1). Forward primers from each microsatellite locus were labeled with fluorescent dyes (Applied Biosystems, Foster City, California, USA and Eurofin MWG, Paris, France): HEX^™^ (for Lv-01), FAM^™^ (for Lv-03 and Lv-16), ATTO565^™^ (for Lv-09), and ATTO550^™^ (for Lv-19). We performed PCR in a 10 µL volume containing 1x multiplex PCR master mix (Qiagen Hilden, Germany), 5-20 ng of genomic DNA, and 0.2 μM of labeled forward and unlabeled reverse primers. The PCR cycling program included an initial denaturation step (95°C for 15 min) followed by 32 cycles of denaturation (94°C for 30 s), annealing (55°C for 1 min 30 s) and extension (72°C for 1 min) and a final extension (60°C for 30 min). We pooled 1.5 µL of the PCR reaction with 0.25 µL of the GeneScan 500 LIZ size standard (Applied Biosystems) and 9.75 µL of deionized formamide. Alleles were size separated by electrophoresis on an ABI PRISM^®^ 3130 sequencer and scored with GeneMapper version 5 (Applied Biosystems). Among the 236 adult trees in the experimental garden, we identified 121 unique genotypes. Each genotype was represented by 2.03 copies on average. These clones derive from plant cuttings that were collected on neighboring individuals *in natura*. Each unique genotype was considered as a potential father in the paternity analysis.

##### Paternity analysis

For the 1209 offspring, paternity was assigned using the maximum-likelihood method described in Kalinowski et al. (2007) and implemented in the program CERVUS version 3.0.7 (Marshall et al. 1998). All unique genotypes in the experimental garden were considered as potential pollen sources. For each offspring-putative father combination, a paternity likelihood was estimated using a ratio of probabilities (LOD score, *i*.*e*. the likelihood that the examined plant is the true father divided by the likelihood that the examined plant is not the true father). To decide whether paternity could be assigned to the individual with the highest LOD score, the difference in LOD scores between the two most likely fathers was calculated and compared to a critical difference in LOD scores below which paternity could not be attributed at a 95% level of confidence. This critical value was obtained using simulations, that were performed using the following parameters: 10 000 offspring, 121 candidate fathers, all candidate fathers sampled and 0.01 as the proportion of mistyped loci. For each offspring, paternity analysis produced three alternative outcomes: (i) no assignment because two or more putative fathers were compatible with the offspring, and the difference in LOD scores was too low to attribute paternity to the most-likely father at the chosen confidence level; (ii) paternity was attributed to the mother, allowing us to estimate a selfing rate for each mother; or (iii) paternity was attributed to another tree.

### Conditions for the maintenance of SC within a DSI system

#### Predictions of inbreeding depression and prior selfing consistent with estimated [SC] phenotype frequencies

Van de Paer et al. (2015) developed a phenotypic model to study the coevolution between self-incompatibility and unisexuality. In particular, they studied the conditions for the maintenance of polymorphism with a pollen-part [SC] phenotype and two [SI] phenotypes, under the assumption that seeds produced by selfing suffer from the cost of inbreeding depression *δ* and that a fraction *γ* of ovules are self-fertilized (see Eq. 15 in Van de Paer et al. 2015). The expected frequency of the pollen-part [SC] phenotype when maintained with both [SI] phenotypes is given by

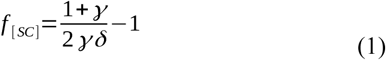

Given the estimated frequency of the [SC] phenotype 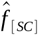 in our set of wild populations and its 95% confidence interval, Eq. (1) was used to determine the sets of values {*δ, γ*} consistent with our observations.

## Results

### *Phenotypic characterization of diallelic SI in* L. vulgare

Among the seven wild populations, stigma tests performed with the pollen from the cross-compatible tester pair detected 94 individuals belonging to the G_A_ SI group and 90 individuals belonging to the G_B_ SI group (Table 1). We did not observe any individuals compatible with both MP-01 and MP-04, thereby excluding the existence of a third incompatibility group among the sampled individuals. Among the 142 individuals tested with their own pollen, 7 (5%) showed an SC reaction with pollen tubes germinating and growing through the stigma and style tissues (Table 1; Fig. 2 Panel C3). These [SC] individuals were detected in five of the seven populations located in geographically and ecologically differentiated regions, from the North Sea coast in France to Bavaria in Germany and from 10 to 1000 m in elevation (Table 1). Strikingly, all the genotypes presenting a [SC] phenotype belonged to the G_A_ group: when tested with pollen from the tester pair, pollen tubes from MP-04 were able to reach the style, but pollen tubes from MP-01 were not. The remaining 135 individuals displayed a typical incompatibility reaction, *i*.*e*. no pollen tubes in the stigma or short pollen tubes that did not reach the style.

### Compatibility/incompatibility relationships among [G_A_], [G_B_], and [SC] individuals assessed using stigma tests in a diallelic design

The stigma tests performed in a diallelic design, where each individual was in turn used as a pollen donor and a pollen recipient, supported the conclusions from the wild population screening regarding the existence of DSI in the study species. The four G_A_ [SI] (CA-44, MP-01, VIF-09, and VIF-16) and four G_B_ [SI] (CA-07, CA-31, G13-08 and MP-04) individuals showed an SI profile (Table 2, Fig. 2 panels A1, B2, D4, and E5) when pollinated with their own pollen. Taken as pollen donors and as pollen recipients, they showed an incompatibility profile when crossed with mates belonging to their own SI group and a compatibility profile when crossed with mates from the other group. For the two [SC] genotypes however the results depended on whether they were used as pollen recipient or as pollen donor. Used as female recipients, the two [SC] individuals showed incompatibility profiles with G_A_ [SI] individuals (Table 2, Fig. 2 panels C1, C2) and compatibility profiles with G_B_ [SI] individuals (Table 2, Fig. 2 panels C4, C5), confirming that they belong to the G_A_ group. Taken as pollen donors, the two [SC] individuals showed a compatibility profile with G_A_ individuals, themselves and G_B_ individuals (Table 2, Fig. 2 column 3).

**Table 2:**
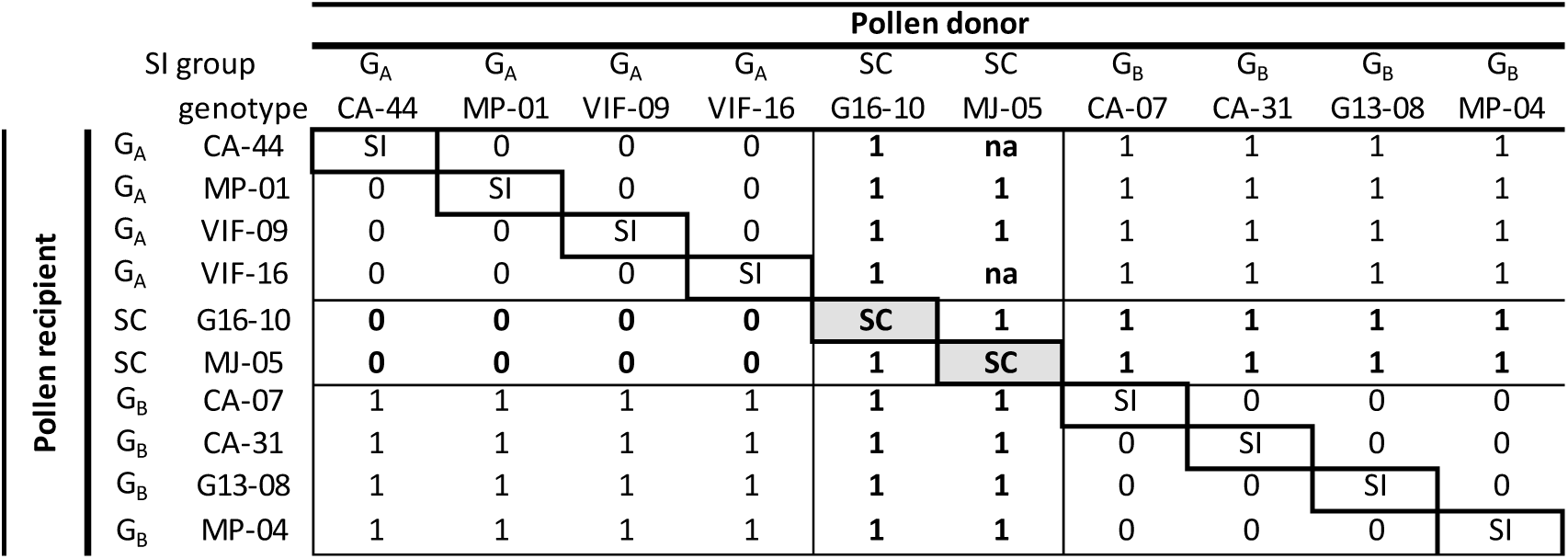
Compatibility relationships between 10 *Ligustrum vulgare* genotypes, characterized at the prezygotic stage using stigma tests in a full diallelic crossing scheme. Germination of pollen tubes on stigmas were scored in 100 crosses (three flowers pollinated in each cross) performed between four G_A_, four G_B_ and two [SC] genotypes (G16-10 and MJ-05). As pollen donors, SC individuals were compatible with both G_A_ and G_B_ genotypes, but as pollen recipients, they were incompatible with G_A_ genotypes and compatible with G_B_ genotypes. SI: self-incompatibility reaction observed (pollen tube growth stopped within stigma); SC: self-compatibility reaction observed (pollen tubes converged in the stigmatic tissue and grew towards the style). 0: incompatibility reaction; 1: compatibility reaction, na: missing data.

|  |  | Pollen donor |  |  |  |  |  |  |  |  |  |
| --- | --- | --- | --- | --- | --- | --- | --- | --- | --- | --- | --- |
|  |  | G<sub>A</sub> | G<sub>A</sub> | G<sub>A</sub> | G<sub>A</sub> | SC | SC | G<sub>B</sub> | G<sub>B</sub> | G<sub>B</sub> | G<sub>B</sub> |
|  |  | CA-44 | MP-01 | VIF-09 | VIF-16 | G16-10 | MJ-05 | CA-07 | CA-31 | G13-08 | MP-04 |
| Pollen recipient | G<sub>A</sub> CA-44 | SI | 0 | 0 | 0 | 1 | na | 1 | 1 | 1 | 1 |
|  | G<sub>A</sub> MP-01 | 0 | SI | 0 | 0 | 1 | 1 | 1 | 1 | 1 | 1 |
|  | G<sub>A</sub> VIF-09 | 0 | 0 | SI | 0 | 1 | 1 | 1 | 1 | 1 | 1 |
|  | G<sub>A</sub> VIF-16 | 0 | 0 | 0 | SI | 1 | na | 1 | 1 | 1 | 1 |
|  | SC G16-10 | 0 | 0 | 0 | 0 | SC | 1 | 1 | 1 | 1 | 1 |
|  | SC MJ-05 | 0 | 0 | 0 | 0 | 1 | SC | 1 | 1 | 1 | 1 |
|  | G<sub>B</sub> CA-07 | 1 | 1 | 1 | 1 | 1 | 1 | SI | 0 | 0 | 0 |
|  | G<sub>B</sub> CA-31 | 1 | 1 | 1 | 1 | 1 | 1 | 0 | SI | 0 | 0 |
|  | G<sub>B</sub> G13-08 | 1 | 1 | 1 | 1 | 1 | 1 | 0 | 0 | SI | 0 |
|  | G<sub>B</sub> MP-04 | 1 | 1 | 1 | 1 | 1 | 1 | 0 | 0 | 0 | SI |

### Compatibility/incompatibility relationships among [G_A_], [G_B_], and [SC] individuals assessed by paternity analysis in the experimental population

All genotyped offspring were compatible with at least one father in the experimental plot. However, we were unable to assign a father to 306 offspring (25%) because two or more individuals were possible fathers with an LOD-score difference too low to attribute paternity to the most likely father at the 95% confidence level.

The overall selfing rate was 0.22 in the experimental population. Among the 201 offspring that were produced by selfing, 200 were collected on the two mother plants that were previously identified as [SC] genotypes on the basis of stigma tests (G16-10 and MJ-05). Although the selfing rates differed between these two genotypes (χ^2^ _(1)_ = 16.77, *P* < 0.001), with values of 0.77 for G16-10 and 0.52 for MJ-05, they were not affected by the stage at which offspring were genotyped (*i*.*e*. embryos *vs*. seedlings, χ^2^ _(1)_ = 7.31.10^−31^, *P* = 1 for G16-10 and χ^2^ _(1)_ = 0.965, *P* = 0.326 for MJ-05). Except for one seedling, outcrossed offspring from the G16-10 [SC] mother were all sired by a tree belonging to the G_B_ group (43 pollination events by five different fathers, see Fig. 3 and Supplementary Table 2). Outcrossed offspring from the other [SC] genotype, MJ-05, were sired either by a G_B_ father (29 pollination events by five different fathers) or by G16-10 (14 pollination events, see Fig. 3 and Supplementary Table 2). Taken as fathers, these two genotypes sired seeds on both G_A_ and G_B_ mothers (see Fig. 3 and Supplementary Table 2). Together, these results confirm the asymmetry that was revealed using the stigma tests: taken as pollen recipients, both [SC] individuals behaved as members of the G_A_ group, but taken as pollen donors, they showed compatibility with both G_A_ and G_B_ individuals.

**Figure 3:**
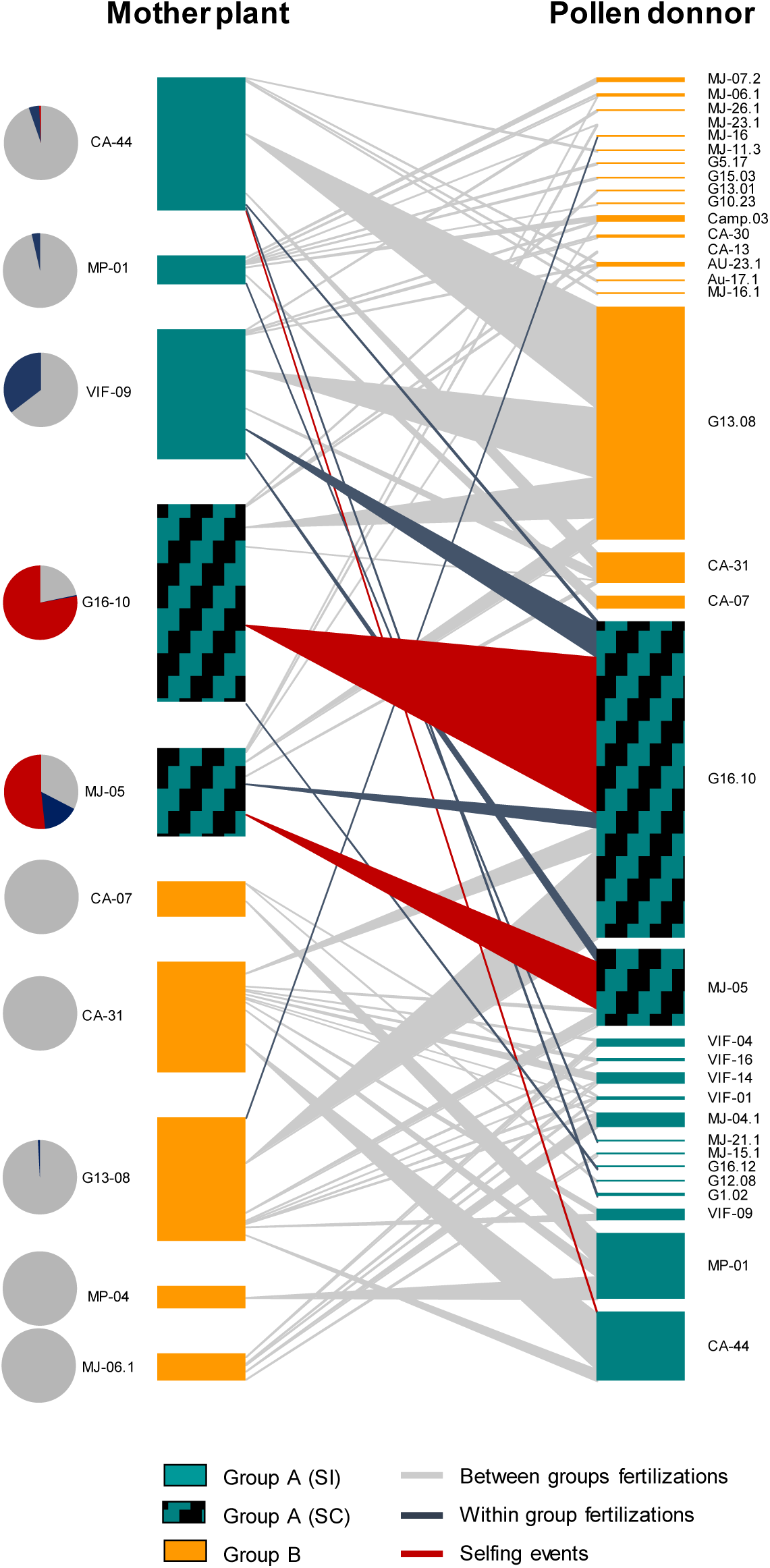
Diagram representing all pollination events detected by paternity analysis in the experimental garden. On the left, each of the 10 mother plants (pollen recipients) is represented by a rectangle whose length is proportional to the number of assigned offspring (from 23 for mother plant MP-04 to 198 for G16-10). On the right, confirmed father plants (pollen donors) are also represented by rectangles whose length is proportional to the number of offspring they sired according to paternity analysis (1 to 315). The width of the links between pollen donors and mother plants are again proportional to the number of detected events and are colored according to the type of event that occurred (*i*.*e*. between-group fertilization, within-group fertilization or selfing event). Pie charts on the left represent the proportion of the different types of cross (between self-incompatibility (SI) groups, within SI group and selfing) detected by paternity assignment for each mother plant.

Regarding pollination patterns in [SI] mothers and potential fathers, 98% of offspring collected on the three G_A_ mothers were sired by G_B_ fathers (236 pollination events by 15 different fathers) and 99% of offspring collected on the five G_B_ mothers were sired by G_A_ fathers (188 pollination events by 10 different fathers, see Fig. 3 and supplementary Table 2), confirming the occurrence of a functional DSI system at the postzygotic stage.

### Conditions for the stability of a polymorphic [SC]/[SI] population in a DSI

The screening for [SC] phenotypes in *L. vulgare* wild populations revealed self-compatible variants occurring at limited frequencies in five out of the seven studied populations (see Table 1). We used the phenotypic model from Van de Paer et al. (2015) to determine the possible range of values of inbreeding depression consistent with a stable polymorphism where both [SI] phenotypes as well as the [SC] phenotype would be maintained in the population with frequencies reflecting the observed values and their confidence intervals (Table 1). Figure 4 shows that the observed frequencies are consistent with a strong inbreeding depression (higher than 0.95) depending on the value of the prior selfing rate, which should be also be substantial (higher than 0.9).

**Figure 4:**
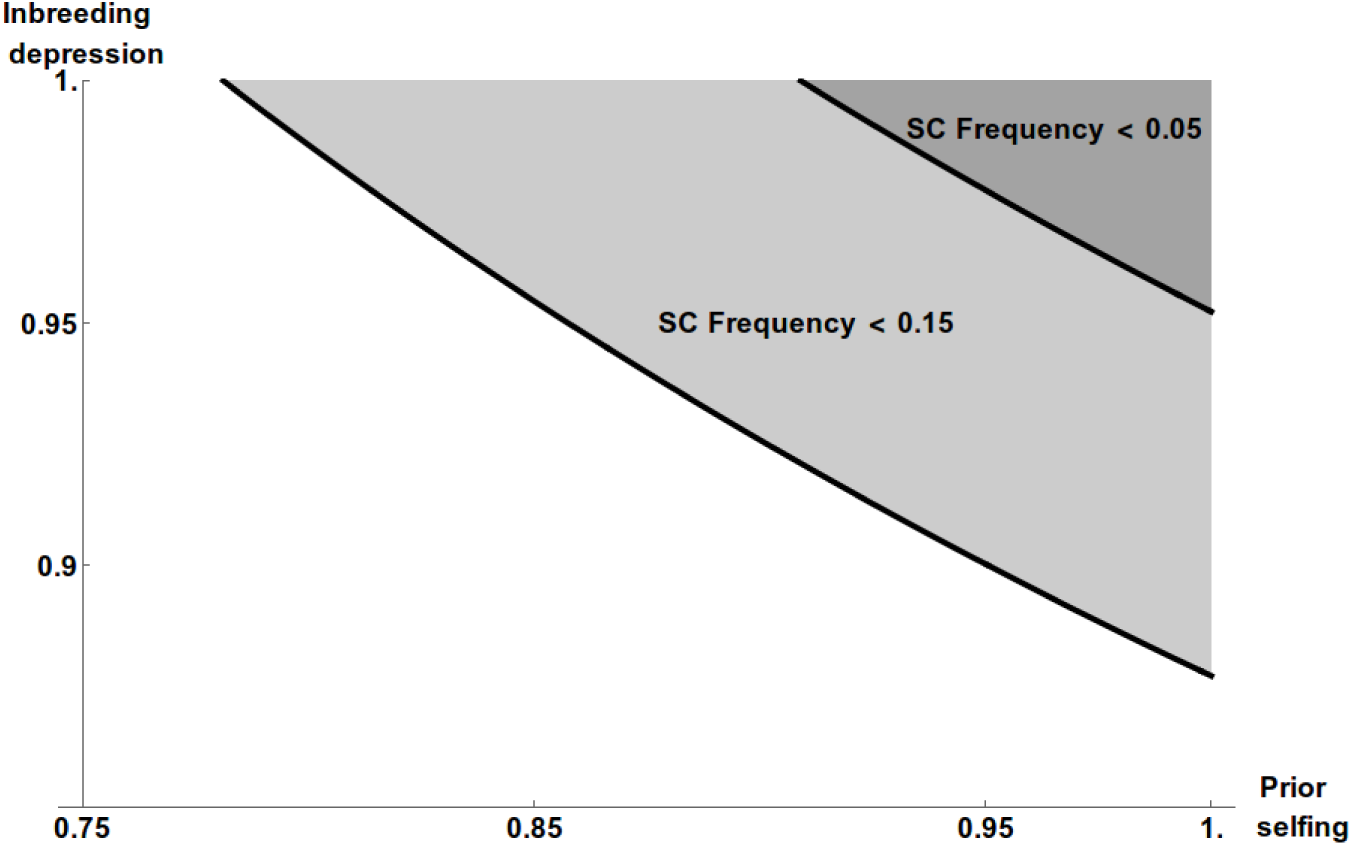
Predicted inbreeding depression and prior selfing rate consistent with the estimated observed frequencies of the self-compatible (SC) phenotype in natural populations of *Ligustrum vulgare*. The curves show the values predicted for the mean (≈ 0.05, top) vs. the lower 95% bound (≈ 0.15, bottom) estimates of the SC phenotype frequency. The gray zone above the curves show the range of values that are consistent with a SC phenotype frequency lower than 0.05 and 0.15. Predictions were obtained using the model built by Van de Paer et al. (2015).

## Discussion

### L. vulgare *possesses DSI with self-compatible variants*

Among all the stigma tests performed during the course of this study, we did not detect a single individual that was compatible with pollen from both G_A_ and G_B_ [SI] testers, confirming the existence of two and only two SI groups within the study populations. *L. vulgare* thus expresses DSI as the other Oleaceae species belonging to the allotetraploid Oleeae tribe that have been tested for SI (Saumitou-Laprade et al. 2010; Vernet et al. 2016; Saumitou-Laprade et al. 2017a; Saumitou-Laprade et al. 2018). According to Oleaceae phylogenetic analyses (Wallander and Albert 2000), the subtribe Ligustrinae forms the basal clade of Oleeae tribe and represents the first lineage having diverged after an allotetraploidization event, before the three other subtribes: Schreberineae (including the genera *Schrebera* and *Comoranthus*), Fraxininae (including the genus *Fraxinus*) and Oleinae (with 12 genera including *Olea* and *Phillyrea*). The existence of two SI groups in *L. vulgare* thus suggests that a homomorphic DSI has been present in the Oleaceae species since the allotetraploidization event. Interestingly, heteromorphic DSI is present in the basal distylous members of the Oleaceae family. Old phylogenetic studies of Oleaceae (Taylor 1945), confirmed by more recent analyses (Wallander and Albert 2000), suggest an ancestral status for the diploid and distylous members of the family. Except in Fontanesieae, heterostyly or distyly is reported in all diploid subtribes. In the Myxopyreae subtribe, *Nyctanthes arbor-tristis* shows heterostyly and pollen dimorphism (Kiew 1984). In the Forsythieae subtribe, the genus *Abeliophyllum* has heterostylous flowers (Ryu 1976) and heterostyly has been known in *Forsythia* since Darwin (Darwin 1877) and described in at least 8 of 11 surveyed species (Kim 1999). In the Jasmineae subtribe, *Jasminum* is reported as hermaphroditic, heterostylous, or usually heterostylous (Woodson et al. 1976; Ganders 1979; Dommée et al. 1992). To date, we do not know whether the self-pollen recognition/rejection capacity expressed in the homomorphic DSI was already present in heterostylous species and has persisted since the breakdown of heterostyly or whether it has evolved following the genomic rearrangements resulting from the allotetraploidization event. Detailed genetic studies and phylogenetic analyses are needed to assess a possible homology of DSI between heterostylous and non-heterostylous taxa in the Oleaceae (Barrett 2019).

### SC behavior in *L. vulgare* involves an SI breakdown in pollen

In multi-allelic SI systems reported in angiosperms, three main model systems are recognized and characterized at the molecular level (Charlesworth et al. 2005). Self-incompatibility is based on recognition of pollen by pistils if both express components related to the same SI specificity. For the sporophytic model in Brassicaceae and the gametophytic models in the Solanaceae and Rosaceae, SI specificity is controlled by a single locus (the S-locus) that contains two closely linked genes expressed in pollen grains and in the style respectively (Takayama and Isogai 2005). Producing a new SI specificity requires changes in both genes, because any mutation affecting only one of the two genes will produce a self-compatible haplotype (Charlesworth and Charlesworth 1979; Uyenoyama et al. 2001; Chantreau et al. 2019). In the present study, prezygotic analyses based on stigma tests and postzygotic analyses based on paternity assignments revealed an asymmetrical compatibility behavior of [SC] individuals depending on whether they were considered as pollen recipients or as pollen donors. As pollen donors, [SC] individuals were compatible with both SI groups whereas, as pollen recipients, they still expressed incompatibility with pollen produced by G_A_ plants. Therefore, the SI breakdown in *L. vulgare* is related to a breakdown in pollen incompatibility, at least in the two [SC] genotypes in our experimental population. In addition, although our wild population screening only examined individuals as pollen recipients (using the tester pair pollen and their own pollen), the seven detected [SC] individuals also behaved as members of the G_A_ group by expressing incompatibility towards pollen produced by the G_A_ tester.

SI breakdown has been observed in other systems, with either breakdown in pollen-part incompatibility (Tsukamoto et al. 2003; Halász et al. 2007; Li et al. 2009) or in style-part incompatibility (e.g. Qi et al. 2011). Although the structure of the DSI locus in Oleaceae family remains unknown, it is clear that the [SC] variants we identified result from a modification in the specificity expressed by pollen produced by plants belonging to the G_A_ group. An open question is whether this modification is due to mutations in a specific S-allele, as reported in species with multi-allelic gametophytic SI systems (Matton et al. 1999; Sonneveld et al. 2005; Wu et al. 2011) and sporophytic SI systems (Chookajorn et al. 2004; Tsuchimatsu et al. 2012) or to the epistatic effect of a modifying factor unlinked to the S-locus but affecting the expression of the pollen gene from a specific S-allele (Wu et al. 2011).

### Evolution and maintenance of the [SI]/[SC] polymorphism

The conditions for the maintenance of SI is a long-standing question in evolutionary biology and has received much theoretical attention (e.g. Charlesworth and Charlesworth 1979, Porcher and Lande 2005b, Gervais et al. 2011, Gervais et al. 2014, Van de Paer et al. 2015). Although evolutionary models predict that the stable coexistence of [SC] and [SI] individuals in a single population is possible, empirical support of such SI/SC polymorphism remains limited. There is evidence that different populations of the same species can be fixed for either [SI] or [SC] in *L. alabamica* (Lloyd 1965) and one population with equifrequent [SI] and [SC] individuals was observed in *A. lyrata* (Mable et al. 2017). The scarcity of polymorphic situations is perhaps not surprising because most self-incompatible species retain large numbers of S-alleles owing to high levels of negative frequency-dependent selection (Castric and Vekemans 2007). When the number of [SI] alleles is locally high, any [SC] variant arising in a population would only have a negligible advantage as compared to the resident [SI] individuals. With DSI and a pollen-part [SC] phenotype, as we now know is the case for *L. vulgare*, the maintenance of such a polymorphic mating system is possible under a range of conditions as long as the number of SI alleles remains low and inbreeding depression is high (Van de Paer et al. 2015). The stability of such polymorphism should rapidly shrink when the number of [SI] groups increases (Van de Paer et al. 2015). This leads to the question whether other hermaphroditic Oleaceae species, such as the olive tree, also show a polymorphic mating system as predicted by theoretical models.

Interestingly, the [SC] variants we observed all belonged to the same SI group and all displayed a behavior that was congruent with the breakdown of pollen-part incompatibility. These variants occurred in populations that were several hundreds of kilometers apart. Two scenarios could explain these observations: (i) either the pollen-part locus in the G_A_ group is somehow more susceptible to breakdown or (ii) [SC] individuals all trace back to the same mutation and it is just a matter of chance that just one incompatibility group was affected and not the other. Whatever the origin of the mutation, theoretical models predict that the polymorphic mating system observed in *L. vulgare* should be maintained only under high levels of inbreeding depression (Fig. 4). However, the fact that (i) germination rates in seeds derived from [SI] and [SC] plants were similar and that (ii) paternity analyses did not reveal differences in selfing rates between the embryo and the seedling stage in [SC] offspring suggest a limited inbreeding depression effect at the germination stage. A possible explanation for the maintenance of the polymorphic mating system observed in *L. vulgare* is that strong inbreeding depression is expressed either at earlier stages of the life cycle (before germination, with seed abortions for instance), or later on, with reduced survival and/or reproduction after the seedling stage. This remains to be estimated since, as shown recently, understanding the evolution of mating systems in perennials requires correctly estimating the lifetime inbreeding depression, and not only what happens during the early stages (Lesaffre and Billiard 2019).

Finally, theoretical models also suggest that SI breakdown in pollen (or in pistils) may represent an evolutionary pathway towards a third SI group through SC intermediates if a compensatory pistil- (or pollen-) part mutation restores SI (Uyenoyama et al. 2001; Gervais et al. 2011; Van de Paer et al. 2015). Interestingly, SI breakdown in pollen may also represent the first step in the evolution towards androdioecy (Van de Paer et al. 2015). The transition from hermaphroditism to stable androdioecy requires at least two mutations: one producing female sterility and one rendering males compatible with both SI groups. Theory helps predict in which order these mutations should appear so as to be selected for. Under nuclear control of sex, models predict that male individuals are eliminated from hermaphrodite populations due to the strong compensation required *via* a fitness advantage for the loss of one sexual function (Lewis 1941; Lloyd 1975; Charlesworth and Charlesworth 1978). It is thus unlikely that female sterility appears first. Additionally, if DSI allows the maintenance of stable androdioecy because males are compatible with both groups of hermaphrodites (Pannell and Korbecka 2010), androdioecy itself facilitates the maintenance of DSI (Van de Paer et al. 2015). The occurrence of compatible males that cannot self and thereby avoid the effects of inbreeding depression prevents the invasion of a SC mutation in the population. Thus, when males are present, [SC] hermaphrodites not only suffer from inbreeding depression, but also from the direct competition with males to sire the SI hermaphrodites (Van de Paer et al. 2015). Therefore, it is more parsimonious to hypothesize that the [SC] mutant appears first, followed by a female sterility mutation closely linked to the SC mutation. In that sense, the SI breakdown in *L. vulgaris* may represent the first step in the evolution towards androdioecy in Oleaceae. Whether the same evolutionary scenario gave rise to the androdioecy documented in other Oleaceae species such as *P. angustifolia* (Saumitou-Laprade et al. 2010) and *F. ornus* (Vernet et al. 2016) remains an open question.

## Author Contributions

All authors contributed to the study presented in this paper. IDC, PV, SB and PS-L. wrote the manuscript. PS-L and PV developed, designed and oversaw the study; they coordinated the collection of data and plant material, carried out stigma tests, participated in data analysis and interpreted the results. IDC performed paternity assignments and data analysis. SB explored the conditions for maintenance of self-compatible mutants at low frequencies. CG developed the molecular markers and performed DNA extractions and genotyping, AB transferred the genotypes from wild populations to the experimental garden at the University of Lille. CP supervised the germination of seeds used for paternity testing, surveyed early seedling growth and collected leaves for DNA extraction.

## Acknowledgements

We thank Nathalie Faure, Eric Schmitt, and Cédric Glorieux for support in caring for the plants at the University of Lille greenhouse. We are very grateful to Géraldine Coste and Sandrine Descave from the Cevennes Natural Park for facilitating access to trees in natural sites at the optimal time in regard to their phenology. We thank John Pannell, Vincent Castric, and Xavier Vekemans for scientific discussions and helpful comments on the manuscript. This research is a contribution to the CPER research project CLIMIBIO. The authors thank the French Ministry for Higher Education and Research, the Hauts-de-France Regional Council and the European Regional Development Fund for their financial support for this project.

## References

Barrett, S. C. 2019. ‘A most complex marriage arrangement’: recent advances on heterostyly and unresolved questions. New Phytologist 224:1051–1067.

Besnard, G., P.-O. Cheptou, M. Debbaoui, P. Lafont, B. Hugueny, J. Dupin, and D. Baali-Cherif. 2020. Paternity tests support a diallelic self-incompatibility system in a wild olive (*Olea europaea subsp. laperrinei,* Oleaceae). Ecology and Evolution 10:1876–1888.

Billiard, S., M. López-Villavicencio, B. Devier, M. E. Hood, C. Fairhead, and Giraud. 2011. Having sex, yes, but with whom? Inferences from fungi on the evolution of anisogamy and mating types. Biological Reviews 86:421–442.

Busch, J. W. and D. J. Schoen. 2008. The evolution of self-incompatibility when mates are limiting. Trends in Plant Science 13:128–136.

Castric, V. and X. Vekemans. 2004. INVITED REVIEW: Plant self-incompatibility in natural populations: a critical assessment of recent theoretical and empirical advances. Molecular Ecology 13:2873–2889.

Castric, V. and X. Vekemans. 2007. Evolution under strong balancing selection: how many codons determine specificity at the female self-incompatibility gene SRK in Brassicaceae? BMC Evolutionary Biology 7:132.

Chantreau, M., C. Poux, M. F. Lensink, G. Brysbaert, X. Vekemans, and V. Castric. 2019. Asymmetrical diversification of the receptor-ligand interaction controlling self-incompatibility in Arabidopsis. eLife 8.

Charlesworth, B. and D. Charlesworth. 1978. A model for the evolution of dioecy and gynodioecy. The American Naturalist 112:975–997.

Charlesworth, D. and B. Charlesworth. 1979. The evolution and breakdown of S-allele systems. Heredity 43:41–55.

Charlesworth, D., X. Vekemans, V. Castric, and S. Glémin. 2005. Plant self-incompatibility systems: a molecular evolutionary perspective. New Phytologist 168:61–69.

Chookajorn, T., A. Kachroo, D. R. Ripoll, A. G. Clark, and J. B. Nasrallah. 2004. Specificity determinants and diversification of the Brassica self-incompatibility pollen ligand. Proceedings of the National Academy of Sciences 101:911–917.

Darwin, C. 1877. The different forms of flowers on plants of the same species. The University of Chicago Press

Dommée, B., J. D. Thompson, and F. Cristini. 1992. Distylie chez *Jasminum fruticans L*.: hypothèse de la pollinisation optimale basée sur les variations de l’écologie intraflorale. Bulletin de la Société Botanique de France. Lettres Botaniques 139:223–234.

Fisher, R. A. 1941. Average excess and average effect of a gene substitution. Annals of Eugenics 11:53–63.

Ganders, F. R. 1979. The biology of heterostyly. New Zealand Journal of Botany 17:607–635.

Gervais, C. E., D. A. Awad, D. Roze, V. Castric, and S. Billiard. 2014. Genetic architecture of inbreeding depression and the maintenance of gametophytic self-incompatibility. Evolution 68:3317–3324.

Gervais, C. E., V. Castric, A. Ressayre, and S. Billiard. 2011. Origin and diversification dynamics of self-incompatibility haplotypes. Genetics 188:625–636.

Goldberg, E. E., J. R. Kohn, R. Lande, K. A. Robertson, S. A. Smith, and B. Igic. 2010. Species selection maintains self-incompatibility. Science 330:493–495.

Goodwillie, C. 1999. Multiple origins of self-compatibility in *Linanthus* section *Leptosiphon* (Polemoniaceae): phylogenetic evidence from internal-transcribed-spacer sequence data. Evolution 53:1387–1395.

Gould, S. J. and E. S. Vrba. 1982. Exaptation. A missing term in the science of form Paleobiology 8:4–15.

Halász, J., A. Pedryc, and A. Hegedűs. 2007. Origin and dissemination of the pollen-part mutated SC haplotype which confers self-compatibility in apricot (*Prunus armeniaca*). New Phytologist 176:792–803.

Igic, B., R. Lande, and J. R. Kohn. 2008. Loss of self-incompatibility and its evolutionary consequences. International Journal of Plant Sciences 169:93–104.

Kalinowski, S. T., M. L. Taper, and T. V. Marshall. 2007. Revising how the computer program CERVUS accommodates genotyping error increases success in paternity assignment. Molecular ecology 16:1099–1106.

Kiew, R. 1984. Preliminary pollen study of the Oleaceae in Malesia. Gardens’ bulletin, Singapore.

Kim, K.-J. 1999. Molecular phylogeny of *Forsythia (Oleaceae)* based on chloroplast DNA variation. Plant Systematics and Evolution 218:113–123.

Lesaffre, T. and S. Billiard. 2019. The joint evolution of lifespan and self-fertilization. Journal of Evolutionary Biology 33:41–56.

Lewis, D. 1941. Male sterility in natural populations of hermaphrodite plants the equilibrium between females and hermaphrodites to be expected with different types of inheritance. New Phytologist 40:56–63.

Li, M., X. Li, Z. H. Han, H. Shu, and T. Z. Li. 2009. Molecular analysis of two Chinese pear (*Pyrus bretschneideri* Rehd.) spontaneous self-compatible mutants, Yan Zhuang and Jin Zhui. Plant Biology 11:774–783.

Lloyd, D. G. 1965. Evolution of self-compatibility and racial differentiation in Leavenworthia (Cruciferae). Contributions from the Gray Herbarium of Harvard University:3–134.

Lloyd, D. G. 1975. The maintenance of gynodioecy and androdioecy in angiosperms. Genetica 45:325–339.

Mable, B. K., J. Hagmann, S. T. Kim, A. Adam, E. Kilbride, D. Weigel, and M. Stift. 2017. What causes mating system shifts in plants? Arabidopsis lyrata as a case study. Heredity 118:52.

Marshall, T. C., J. Slate, L. E. B. Kruuk, and J. M. Pemberton. 1998. Statistical confidence for likelihood-based paternity inference in natural populations. Molecular Ecology 7:639–655.

Matton, D. P., D. T. Luu, Q. Xike, G. Laublin, M. O’Brien, O. Maes, D. Morse, and M. Cappadocia. 1999. Production of an S RNase with dual specificity suggests a novel hypothesis for the generation of new S alleles. The Plant Cell 11:2087–2097.

Obeso, J. and P. Grubb. 1993. Fruit maturation in the shrub *Ligustrum vulgare (Oleaceae*): lack of defoliation effects. Oikos:309–316.

Pannell, J. R. and G. Korbecka. 2010. Mating-system evolution: rise of the irresistible males. Current Biology 20:R482–R484.

Porcher, E. and R. Lande. 2005a. The evolution of self-fertilization and inbreeding depression under pollen discounting and pollen limitation. Journal of evolutionary biology 18:497–508.

Porcher, E. and R. Lande. 2005b. Loss of gametophytic self-incompatibility with evolution of inbreeding depression. Evolution 59:46–60.

Qi, Y.-J., Y.-T. Wang, Y.-X. Han, S. Qiang, J. Wu, S.-T. Tao, S.-L. Zhang, and H.-Q. Wu. 2011. Self-compatibility of ‘Zaoguan’(*Pyrus bretschneideri* Rehd.) is associated with style-part mutations. Genetica 139:1149–1158.

Ryu, T. 1976. Studies on heterostyly incompatibility of *Abeliophyllum distichum*. Seoul Nat. Univ., Coll. Agri. Bull. 1:113–120.

Saumitou-Laprade, P., P. Vernet, A. Dowkiw, S. Bertrand, S. Billiard, B. Albert, P.-H. Gouyon, and M. Dufay. 2018. Polygamy or subdioecy? The impact of diallelic self-incompatibility on the sexual system in *Fraxinus excelsior* (Oleaceae). Proceedings of the Royal Society B: Biological Sciences 285.

Saumitou-Laprade, P., P. Vernet, C. Vassiliadis, Y. Hoareau, G. de Magny, B. Dommée, and J. Lepart. 2010. A self-incompatibility system explains high male frequencies in an androdioecious plant. Science 327:1648–1650.

Saumitou-Laprade, P., P. Vernet, X. Vekemans, S. Billiard, S. Gallina, L. Essalouh, A. Mhaïs, A. Moukhli, A. Bakkali, G. Barcaccia, F. Alagna, R. Mariotti, N. G. M. Cultrera, S. Pandolfi, M. Rossi, B. Khadari, and L. Baldoni. 2017a. Elucidation of the genetic architecture of self-incompatibility in olive: Evolutionary consequences and perspectives for orchard management. Evolutionary Applications 10:860–866.

Saumitou-Laprade, P., P. Vernet, X. Vekemans, V. Castric, G. Barcaccia, B. Khadari, and L. Baldoni. 2017b. Controlling for genetic identity of varieties, pollen contamination and stigma receptivity is essential to characterize the self-incompatibility system of *Olea europaea L*. Evolutionary Applications 10:867–880.

Signorell, A. 2019. DescTools: Tools for descriptive statistics. R package version 0.99.28 17.

Sison, C. P. and J. Glaz. 1995. Simultaneous confidence intervals and sample size determination for multinomial proportions. Journal of the American Statistical Association 90:366–369.

Sonneveld, T., K. R. Tobutt, S. P. Vaughan, and T. P. Robbins. 2005. Loss of pollen-S function in two self-compatible selections of Prunus avium is associated with deletion/mutation of an S haplotype–specific F-Box gene. The Plant Cell 17:37–51.

Stebbins, G. L. 1957. Self fertilization and population variability in the higher plants. The American Naturalist 91:337–354.

Takayama, S. and A. Isogai. 2005. Self-incompatibility in plants. Annu. Rev. Plant Biol. 56:467–489.

Taylor, H. 1945. Cyto-taxonomy and phylogeny of the Oleaceae. Brittonia 5:337–367.

Tsuchimatsu, T., P. Kaiser, C.-L. Yew, J. B. Bachelier, and K. K. Shimizu. 2012. Recent loss of self-incompatibility by degradation of the male component in allotetraploid *Arabidopsis kamchatica*. PLoS Genetics 8:e1002838.

Tsukamoto, T., T. Ando, K. Takahashi, T. Omori, H. Watanabe, H. Kokubun, E. Marchesi, and T.-h. Kao. 2003. Breakdown of self-incompatibility in a natural population of *Petunia axillaris* caused by loss of pollen function. Plant Physiology 131:1903–1912.

Uyenoyama, M. K., Y. Zhang, and E. Newbigin. 2001. On the origin of self-incompatible haplotypes: transition through self-compatible intermediates. Genetics 157:1805–1817.

Van de Paer, C., P. Saumitou-Laprade, P. Vernet, and S. Billiard. 2015. The joint evolution and maintenance of self-incompatibility with gynodioecy or androdioecy. Journal of Theoretical Biology 371:90–101.

Vernet, P., P. Lepercq, S. Billiard, A. Bourceaux, J. Lepart, B. Dommée, and P. Saumitou-Laprade. 2016. Evidence for the long-term maintenance of a rare self-incompatibility system in Oleaceae. New Phytologist 210:1408–1417.

Wallander, E. and V. A. Albert. 2000. Phylogeny and classification of Oleaceae based on *rps16* and *trnL-F* sequence data. American Journal of Botany 87:1827–1841.

Woodson, R. E., R. W. Schery, and W. G. D’Arcy. 1976. Flora of Panama. Part VIII. Family 158. Oleaceae. Annals of the Missouri Botanical Garden 63:553–564.

Wu, J., C. Gu, Y.-H. Du, H.-Q. Wu, W.-S. Liu, N. Liu, J. Lu, and S.-L. Zhang. 2011. Self-compatibility of ‘Katy’apricot (*Prunus armeniaca L*.) is associated with pollen-part mutations. Sexual plant reproduction 24:23–35.

