## Supplemental Figure 1, Table 1 and Table 2 for "Widespread coexistence of self-compatible and self-incompatible phenotypes in a diallelic self-incompatibility system in *Ligustrum vulgare* (Oleaceae)"

**Supplementary Figure 1**. Map of the artificial *Ligustrum vulgare* population established in the experimental garden at the University of Lille (50°36’28.0”N 3°08’37.1”E). A total of 236 ramets representing 121 genotypes were distributed into four plots. *The 10 individual ramets on which seeds were collected to perform paternity analyses: 6 in plot 1, 2 in plot 3 and 1 in plot 4.

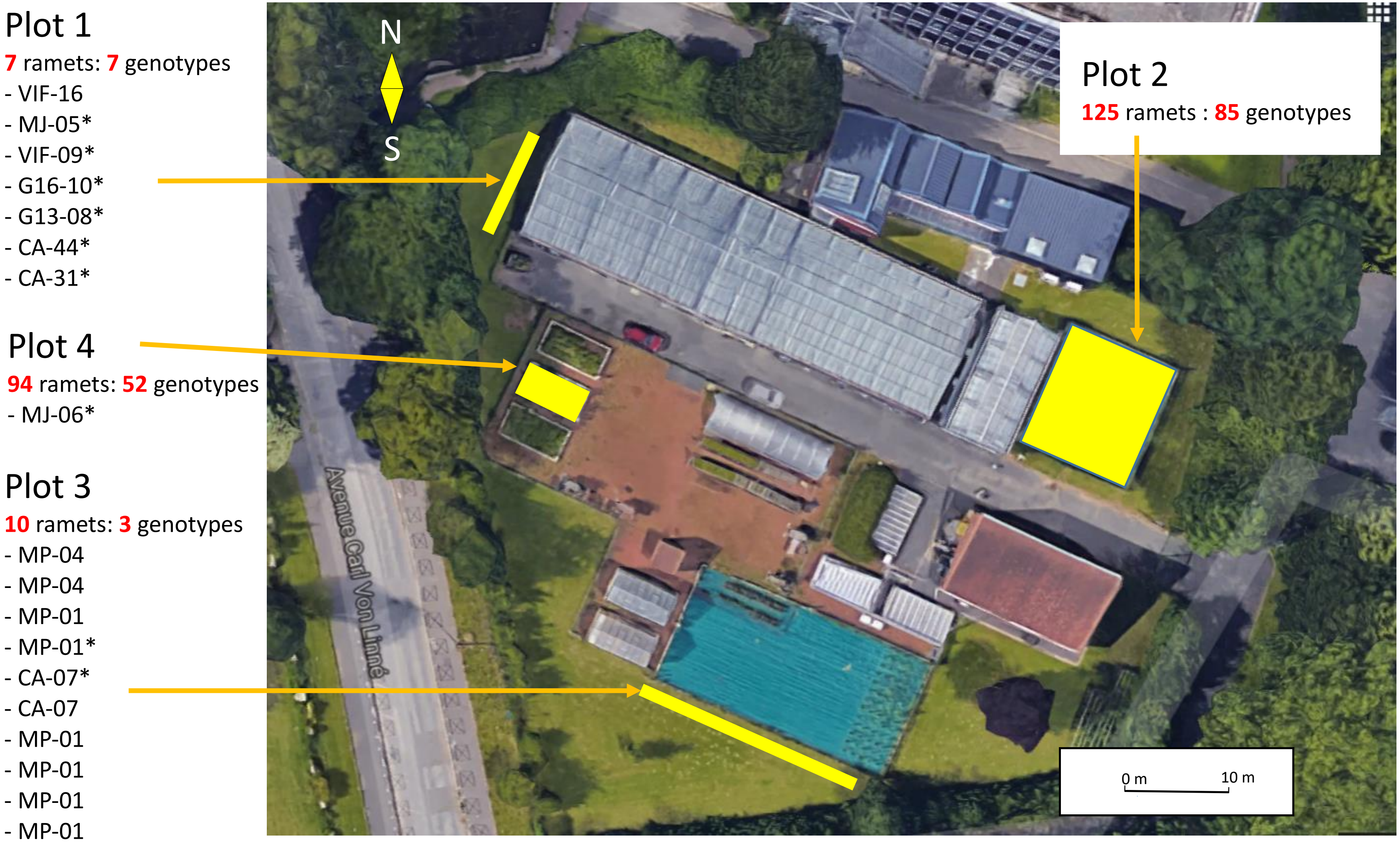

**Supplementary Table 1**: Characteristics of the five polymorphic microsatellite markers developed in *Ligustrum vulgare*. Name, primer sequence (3'-5'), repeat motif from the original sequence, dye used and accession number of six highly polymorphic microsatellite markers isolated in the European privet *Ligustrum vulgare* amplified in a single multiplex PCR reaction. The allele size range (in base pairs) is given for 119 genotypes sampled in four European regions (see detailed description in Table 1). Population 1: North Sea coast (25 genotypes); Population 2: Bavaria (36 genotypes), Population 3: Cevennes (45 genotypes); Population 4: Alps (13 genotypes).

| Locus | Primer sequences (3'-5') | Repeat motif | Dye | GenBank accession n° | Allele size range (bp) | Population 1 | | | Population 2 | | | Population 3 | | | Population 4 | | | All | |
|---|---|---|---|---|---|---|---|---|---|---|---|---|---|---|---|---|---|---|---|
| | | | | | | NA | He | Fis | NA | He | Fis | NA | He | Fis | NA | He | Fis | NA | He |
| Lv-01 | F: TCAATTTCCATGTCAAGACTTCAT<br>R: ACCAACACCACAACAATGATAA | (CA)$_{25}$ | HEX | MN936083 | 221-259 | 12 | 0.818 | 0.134 | 13 | 0.923 | 0.44 | 10 | 0.84 | -0.005 | 11 | 0.923 | 0.167 | 18 | 0.801 |
| Lv-03 | F: TTATCATGGTCCGAATCAACC<br>R: TGGATGATAAATGGAGCCAA | (TTC)22 | FAM | MN936084 | 259-302 | 9 | 0.884 | 0.01 | 9 | 0.798 | 0.115 | 5 | 0.772 | 0.052 | 12 | 0.856 | 0.191 | 14 | 0.752 |
| Lv-09 | F: TGTAACTTCAGCCTTTGCCA<br>R: TCTTGAGTCGTGAGTGTCGG | (GA)18 | ATTO565 | MN936087 | 230-247 | 7 | 0.764 | 0.073 | 7 | 0.829 | 0.078 | 5 | 0.526 | -0.038 | 8 | 0.798 | 0.229 | 9 | 0.658 |
| Lv-16 | F: GCTGTTACAATCCTACCCCTT<br>R: TCTTATGGCGAAGTACCGCT | (TTC)17 | FAM | MN936088 | 136-193 | 14 | 0.886 | 0.153 | 11 | 0.913 | 0.0384 | 6 | 0.744 | 0.236 | 11 | 0.881 | 0.04 | 19 | 0.682 |
| Lv-19 | F: CAACCAAACAGATTAAATACACATACA<br>R: TTGGCAAGGTACACATCTGG | (AC)16 | ATTO550 | MN936089 | 200-215 | 5 | 0.685 | -0.034 | 6 | 0.719 | -0.146 | 3 | 0.253 | 0.034 | 4 | 0.686 | 0.215 | 8 | 0.579 |
| All | | | | | | | 0.0542 | ns | | 0.0208 | ns | | 0.0438 | ns | | 0.0063 | ns | | |

NA, allele number; He, expected heterozygosity; Fis,within population inbreeding coefficient (all values show non significant deviation of Hardy Weinberg equilibriumbased on randomisations)

**Supplementary Table 2**: Cross-compatible and selfing events assessed at the postzygotic stage by paternity analysis in 10 open pollinated *Ligustrum vulgare* progenies. In italics, the progenies resulting either from within-self-incompatibility (SI)-group crosses or from selfing. *Progenies assigned to different [$G_A$] fathers (10 individuals) present in the experimental garden. **Progenies assigned to different [$G_B$] fathers (15 individuals) present in the experimental garden. ***Paternity assigned to a father which phenotype was not tested.

| | | | | **Father characterized fo SI phenotype identified within the artificial population** | | | | | | | | | | | | | **Total number of sampled progenies** | | | | | | | | % selfing in assigned samples | | |
|---|---|---|---|---|---|---|---|---|---|---|---|---|---|---|---|---|---|---|---|---|---|---|---|---|---|---|---|
| | | | | CA-44 | MP-01 | VIF-09 | *10 $G_A$ Fathers | G16-10 | MJ-05 | CA-31 | CA-07 | G13-8 | MJ-06.1 | MP-04 | **15 $G_B$ fathers | *** 7 not phenotyped fathers | genotyped | | assigned to a father (95) | | assigned to a phenotyped father 95 | | produced by selfing | | | | |
| **Open pollinated mother** | **SI Phenotype** | **genotype** | **Position** | $G_A$ | | | | SC | | $G_B$ | | | | | | | Embryo | seedling | Embryo | seedling | Embryo | seedling | Embryo | seedling | Embryo | seedling | Overall |
| | $G_A$ | CA-44 | Plot-1_6 | ***1*** | *0* | *0* | *3* | *3* | *0* | 19 | 0 | 102 | 0 | 0 | 4 | 1 | 50 | 122 | 41 | 92 | 41 | 91 | 0 | 1 | 0.0 | 1.1 | 0.8 |
| | | MP-01 | Plot-3_4 | *0* | *0* | *0* | *1* | *0* | *0* | 0 | 13 | 0 | 2 | 0 | 2 | 1 | 50 | - | 29 | - | 28 | - | 0 | - | 0.0 | - | 0.0 |
| | | VIF-09 | Plot-1_3 | *0* | *0* | *0* | *0* | *34* | *12* | 9 | 0 | 68 | 0 | 0 | 7 | 1 | 50 | 111 | 40 | 91 | 40 | 90 | 0 | 0 | 0.0 | 0.0 | 0.0 |
| | SC | G16-10 | Plot-1_4 | *0* | *0* | *0* | *1* | ***154*** | *0* | 1 | 0 | 39 | 0 | 0 | 3 | 3 | 50 | 201 | 43 | 158 | 43 | 155 | 33 | 121 | 76.7 | 76.6 | 76.6 |
| | | MJ-05 | Plot-1_2 | *0* | *0* | *0* | *0* | *14* | ***46*** | 3 | 0 | 23 | 1 | 0 | 2 | 0 | 50 | 53 | 41 | 48 | 41 | 48 | 24 | 22 | 58.5 | 45.8 | 51.7 |
| | $G_B$ | CA-31 | Plot-1_7 | 56 | 11 | 5 | 11 | 22 | 4 | *0* | *0* | *0* | *0* | *0* | *0* | 2 | 50 | 103 | 42 | 69 | 41 | 68 | 0 | 0 | 0.0 | 0.0 | 0.0 |
| | | CA-07 | Plot-3_5 | 0 | 33 | 0 | 2 | 0 | 0 | *0* | *0* | *0* | *0* | *0* | *0* | 0 | 50 | - | 35 | - | 35 | - | 0 | - | 0.0 | - | 0.0 |
| | | G13-8 | Plot-1_5 | 11 | 0 | 6 | 4 | 88 | 14 | *0* | *0* | *0* | *0* | *0* | *1* | 1 | 50 | 112 | 36 | 89 | 36 | 88 | 0 | 0 | 0.0 | 0.0 | 0.0 |
| | | MJ-06.1 | Plot-4_L3 | 0 | 0 | 0 | 26 | 0 | 0 | *0* | *0* | *0* | *0* | *0* | *0* | 0 | 50 | - | 26 | - | 26 | - | 0 | - | 0.0 | - | 0.0 |
| | | MP-04 | Plot-3_2 | 0 | 23 | 0 | 0 | 0 | 0 | *0* | *0* | *0* | *0* | *0* | *0* | 0 | 48 | - | 23 | - | 23 | - | 0 | - | 0.0 | - | 0.0 |
| | | Total | | 68 | 67 | 11 | 48 | 315 | 76 | 32 | 13 | 232 | 3 | 0 | 19 | 9 | 498 | 702 | 356 | 547 | 354 | 540 | 57 | 144 | | | |
